# Targeting the host protein G3BP1 for the discovery of novel antiviral inhibitors against Chikungunya virus

**DOI:** 10.1101/2022.11.11.516135

**Authors:** Supreeti Mahajan, Ravi Kumar, Ankur Singh, Akshay Pareek, Pravindra Kumar, Shailly Tomar

**Affiliations:** Department of Biosciences and Bioengineering, Indian Institute of Technology Roorkee, Roorkee, Uttarakhand, India– 247667

**Author notes:** The authors have contributed equally (Supreeti Mahajan, Ravi Kumar). Address correspondence to Prof. Shailly Tomar, Department of Biosciences and Bioengineering, Indian Institute of Technology Roorkee, Roorkee, Uttarakhand, India – 247667, Email ID of corresponding author.

**Keywords:** Chikungunya, Host protein G3BP1, Stress granules, Antiviral, Drug discovery

## Abstract

Molecular interactions of Chikungunya virus (CHIKV) nsP3 with the host G3BP1 stress granule (SG) protein is crucial for CHIKV replication. NsP3 binds to the nuclear transport factor 2 (NTF2)-like domain of G3BP1 via its two FGDF motifs, unsettling SGs formation. The present study identified seven small molecules targeting the FGDF binding pocket of G3BP1 using structure-based computer-aided drug discovery. All seven molecules exhibited good binding affinities in the range of ∼3.4 to ∼98μM towards NTF2-like G3BP1 domain. Furthermore, identified molecules demonstrate dose-dependent inhibition in CHIKV infected Vero cells (EC_50_: 0.40 to 7.39µM), and reduced SGs formation in these cells. These results highlight G3BP1 protein as a potential CHIKV therapeutic target and offer potential prospective for developing treatment for CHIKV disease.

## Introduction

Viral infections are imminently accompanied by the exploitation of host resources through continual interaction with the host proteins, resulting in an intricate network of virus-host interactions. Viruses possess few genes and limited biosynthetic capability; therefore, they depend on host factors for efficient viral replication [1–2]. Various host antiviral response mechanisms also operate in defence to the early viral infection and its control [3–4]. Consequently, viruses devise counteracting mechanisms against host antiviral responses to establish viral infection and spread. Additionally, viruses exploit host proteins to modify the metabolic pathways involved in cellular defence mechanisms [5–8]. Viral infections inside a host cell can lead to intrinsic stress conditions that induce the formation of cytoplasmic granules, known as stress granules (SGs) [9–12]. SGs are electron-dense cytoplasmic structures that are non-membranous and are defined by the presence of various RNA binding proteins like TIA-1, TTP, TIAR, Ataxin-2, G3BP1, stalled translation initiation complexes and RNA aggregates [13]. Induction of SGs is triggered when translation initiation is inhibited due to cellular stresses including viral infections. Interestingly, several viruses belonging to diverse families have developed various mechanisms to modulate the formation and functioning of SGs. Previous studies have reported that viruses such as Sindbis virus (SINV), Poliovirus, etc can induce the formation of stress granules at some point during their replication cycle which appears to be advantageous for the replication of viruses [14–15]. However, some studies have suggested that the presence of stress granules is detrimental to the virus infection as in case of Semliki Forest virus (SFV), Dengue virus (DENV), and Chikungunya virus (CHIKV) [16–17]. In general, SG assembly appears to be a crucial component of the cellular antiviral response so that its formation is modulated by the viruses for their gene expression and replication [18]. The major player in the assembly of cellular SGs is G3BP protein [12,19], a multifunctional RNA binding protein that exists in three human isoforms: G3BP1, G3BP2a, and G3BP2b. G3BP1 possesses RNA recognition motifs (RRM) which is required for the induction of SGs along with other domains. The N-terminus of G3BP1 consists of a nuclear transport factor 2 (NTF2)-like domain which plays a role in dimerization of protein [20–21].

Interestingly, in the viral infections, G3BP1 protein can play variety of functions. One of the many functions of G3BP1 is to prevent viral replication through various mechanisms such as G3BP1 interacts with specific viral proteins and not able to form SGs efficiently to produce cellular antiviral response. For instance, G3BP1 is sequestered by molecular interaction with specific viral proteins such as nsP3 of Old world alphaviruses (CHIKV/SFV) [22–24]. G3BP1 interacts with the internal ribosome entry site (IRES) in case of foot-and-mouth disease virus (FMDV) resulting in a change in the structure of IRES [25]. In mammalian reovirus (MRV), the non-structural proteins σNS and μNS interact with G3BP1 and inhibits the replication of MRV [26]. SARS-CoV-2 nucleocapsid protein has been proposed to interact with G3BP1 and impairs G3BP1-mediated SG formation [27]. In Human immunodeficiency virus (HIV) infection, Gag protein has been identified to interact with G3BP1, which modulates the SG assembly [28]. Moreover, in the other viral mechanism, G3BP1 is cleaved by the viral proteases, which leads to the manipulation of stress activated SG pathway and innate antiviral response in viral infected cells. For instance, the lead protease (L^pro^) of and the 3C protease (3C^pro^) of FMDV cleave G3BP1 [25,29,30], Human enterovirus D68 (EV-D68) cleaves G3BP1, which leads to the disassembly of SGs [31]. In a few viruses, G3BP1 inhibits viral replication through positively regulating retinoic acid-inducible gene 1 (RIG-1) mediated antiviral response in the infected cells [32] such as in Enterovirus 71 (EV71), Sendai virus (SeV), etc [31]. Thus, viruses have developed their own ways to inhibit the formation of SGs and overcome the antiviral mechanisms exerted by G3BP1 to promote efficient viral genome replication in the infected cells.

Nonetheless, many viruses utilize G3BP1 to promote virus replication and proliferation. These viruses use G3BP1 as a proviral factor because it plays a crucial role in the viral replication cycles. Vaccinia virus (VV), a DNA virus, uses G3BP1 as a proviral factor, which is active at distinct stages of virus replication [33]. In alphaviruses (CHIKV/SFV) also, the G3BP1 host protein has been shown to support viral replication [34,22]. It has been demonstrated in some viruses that G3BP1 is recruited to replication complexes (RCs) or transcriptional complexes. Interestingly, this indirectly blocks SG assembly as G3BP1 is not available. In Old World alphaviruses (CHIKV/SFV), the FGDF motif present in the C-terminus of viral non-structural protein 3 (nsP3) has been proposed to recruit G3BP1 into viral replication complexes, which in turn recruits host translational machinery to the site. The RGG domain of G3BP1 binds to the 40S ribosomal subunit to recruit the host translational machinery for translation of the viral mRNAs [22]. A similar FGDF motif is also present in HSV-type 1 (HSV-1) ICP8 protein and has been shown to inhibit the SG formation after binding to G3BP1 [35]. Thus, FGDF motif binding pocket in G3BP1 is a potential antiviral target site for therapeutic molecular interventions.

CHIKV is a re-emerging arbovirus of the family *Togaviridae*. Since 2005, CHIKV has re-emerged and initiated large outbreaks in Asia, America and Africa [36]. CHIKV infects distinct vertebrate and invertebrate hosts, including humans [37]. It adversely affects the health of humans by causing mild to debilitating disease with symptoms ranging from fever, muscle pain, joint swelling, headache, nausea, fatigue, and severe joint pain, and rash to polyarthralgia that may persist for days to weeks [38]. The CHIKV genome constitutes two open reading frames (ORFs)- ORF1 and ORF2 embedded between the 5′ and 3′ untranslated regions (UTRs). ORF1 encodes a non-structural polyprotein (P1234) which is further cleaved into four proteins - nsP1, nsP2, nsP3, and nsP4. These nsPs are necessary for the viral genome replication and translation to form replicase complex inside the host cell [39–43]. nsP3 of CHIKV has been shown to interact with the host factor G3BP1 through two tandem FGDF motifs at the C-terminus of the protein [23]. nsP3 is believed to function by disrupting the SGs and RNA/protein subcellular structures formed during viral infections [44,45]. nsP3 mediates the recruitment of the host protein G3BP1 via its two tandem FGDF motifs that bind to a hydrophobic groove on the surface of the NTF2 domain of G3BP1 resulting in its sequestration [46,10]. Hence, CHIKV exploits the host SG effect or proteins like G3BP1 to maximize its replication efficiency. Similarly, SFV also blocks SG assembly by targeting G3BP1. When nsP3-G3BP1 interaction is absent, the replication of SFV is compromised yet detectable [35, 44]. Crystal structure of G3BP1 in complex with FGDF peptide derived from SFV nsP3 is available at Protein Data Bank (PDB ID 5FW5) [46]. Presently, there are no licensed antivirals or vaccines available in the market against CHIKV.

Inspired by the observation that CHIKV replication is severely attenuated in the absence of the nsP3-G3BP interaction, this study targets G3BP1 as a potential host antiviral target and identified novel antivirals against CHIKV infection. The three-dimensional structure of the G3BP1-FGDF peptide complex structure provides a detailed understanding of the key molecular interactions. Using *in silico* virtual screening and docking approach, seven potential antiviral compounds that bind to the G3BP1-FGDF interaction pocket have been identified. Biophysical techniques were employed to measure the binding thermodynamics of identified molecules to the target protein. The antiviral potential of G3BP1-binding molecules against the clinical strain of CHIKV has also been assessed. Immunofluorescence assay (IFA) was established to investigate the relationship between CHIKV replication and virus induced SGs. This is the first study that has targeted the key molecular interactions between nsP3 FGDF-motif of CHIKV with the NTF2-like domain of the host protein G3BP1 and small molecules with high antiviral efficacy have been identified.

## Results

### Structure-based identification of G3BP1 binding molecules

Virtual screening of the LOPAC^1280^ library was done against the FGDF binding pocket in the G3BP1 NTF2-like domain using AutoDock Vina in PyRx 0.8. The binding energy (B.E.) of screened compounds was compared with that of G3BP1-FGDF peptide complex, and protein-ligand complexes with highest B.E. and the best-fit were selected for molecular docking studies. Docking studies were performed for the top seven compounds L-7, WIN, SB2, NAL, DHD, GSK, and FLU into the FGDF motif binding pocket in the G3BP1 NTF2-like domain. The compounds exhibited B.E values in the range of −12.65 to −9.85 kcal/mol, which was considerably higher than that of FGDF (−8.08 kcal/mol) (**Fig.1**) (**Table 1**).

**Fig.1.**
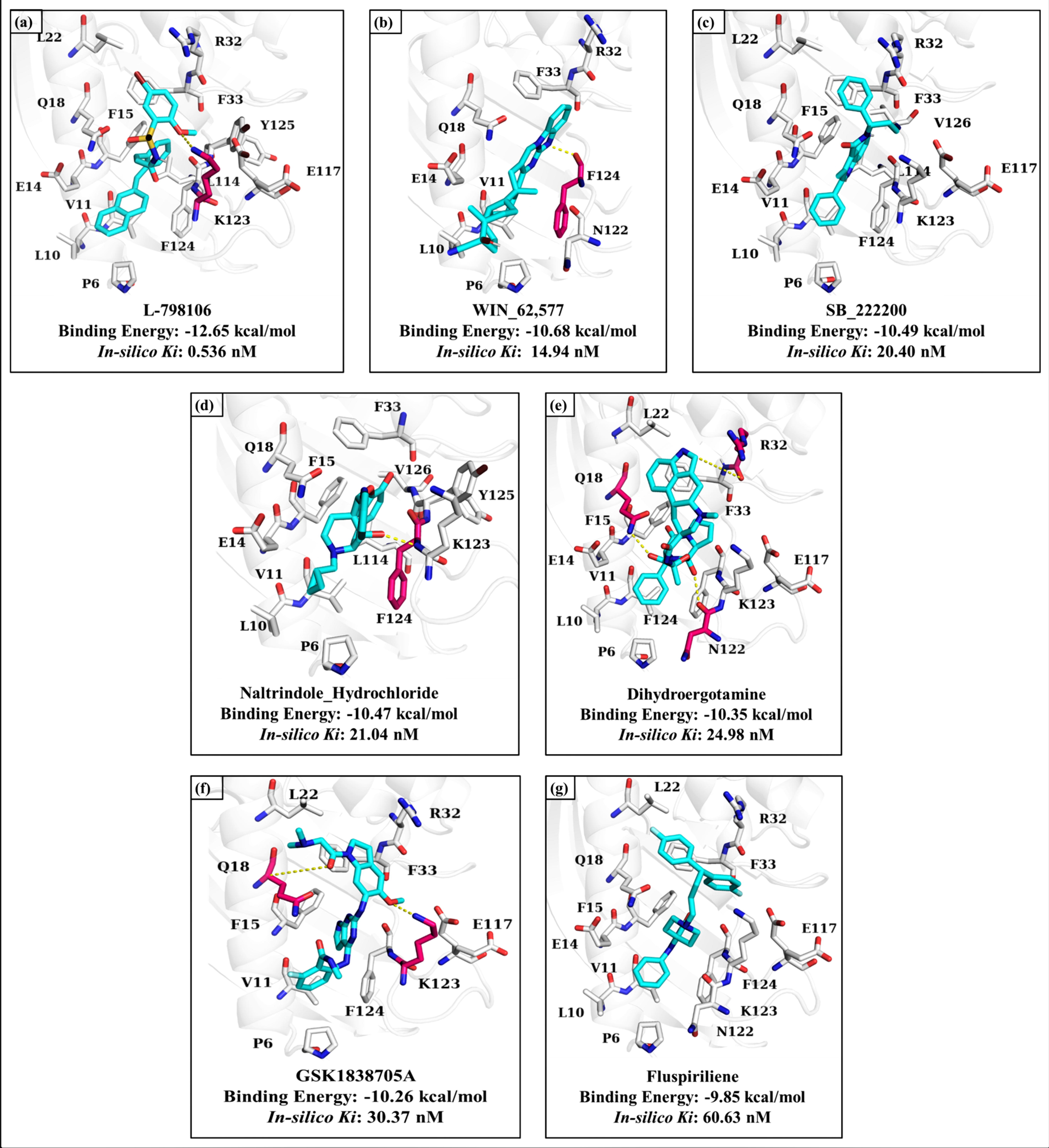
Schematic representation of putative binding mode of compounds in the FGDF motif binding pocket of G3BP1 with. (a) L-7, (b) WIN, (c) SB2, (d) NAL, (e) DHD, (f) GSK, and (g) FLU complexes. Compounds are shown in cyan coloured sticks, residues of G3BP1 with H-bond interactions are depicted in magenta-coloured sticks, and the residues with hydrophobic interactions are shown in white colored sticks.

**Table 1.** Binding energies and *in-silico* inhibition constant (*K_i_*) obtained after completion of docking analysis of screened compounds with G3BP1.

| Compounds | *in silico* inhibition Constant ($K_i$) | Binding Energy (kcal/mol) |
| --- | --- | --- |
| L-798106 | 0.536 nM | -12.65 |
| WIN-62577 | 14.94 nM | -10.68 |
| SB_222200 | 20.40 nM | -10.49 |
| Naltrindole Hydrochloride | 21.04 nM | -10.47 |
| Dihydroergotamine | 24.98 nM | -10.35 |
| GSK1838705A | 30.37 nM | -10.26 |
| Fluspiriline | 60.63 nM | -9.85 |

Using PyMol and Ligplot^+^, the hydrogen bond/hydrophobic interactions in the best fit conformations of the selected compounds within the FGDF binding pocket of G3BP1 protein were analysed (**Fig.1** and **Fig.2**). Detailed analysis of molecular interactions showed that DHD makes three hydrogen bonds (H-bond), and GSK interacts with two H- bonds, whereas L-7, WIN and NAL make only one H- bond each. These seven compounds have two common residues for H-bond interactions *i.e.* Lys123 and Phe124 and also common residues that make hydrophobic interactions with the ligand molecules (**Fig.2**). The common molecular interactions at the protein-ligand interface suggest that the seven compounds may be effective as inhibitors of the CHIKV nsP3-G3BP1 interaction and thus virus replication inhibitors. Additionally, these data provide structure-based hints toward design, modification, and development of more effective analogues of these identified molecules.

**Fig.2.**
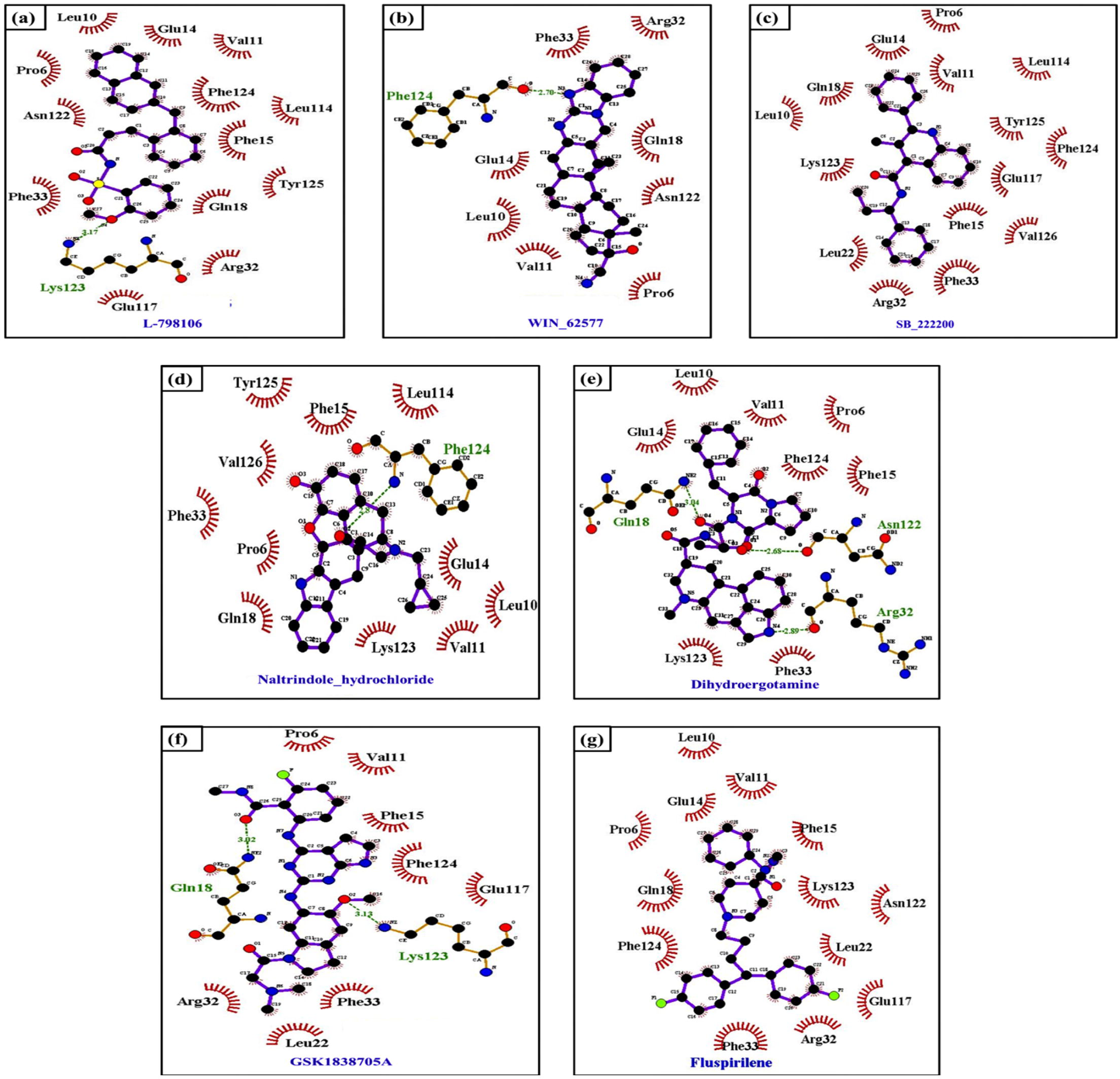
Two-dimensional pictorial representation of interaction involved in binding of inhibitor(s) to the FGDF motif binding pocket of G3BP1. (a) L-7, (b) WIN, (c) SB2, (d) NAL, (e) DHD, (f) GSK, and (g) FLU. Ligands are represented in purple color, H- bonds are shown in green dotted lines, residues involved in H-bond interactions are displayed in green colour, red stellation represents hydrophobic interactions. The two-dimensional figures were prepared using LigPlot+

Molecular dynamic simulation studies of G3BP1 in complex with identified molecules were performed. Analysis of various molecular dynamic parameters- root mean square deviation (RMSD), root mean square fluctuation (RMSF), Radius of gyration (Rg) and H-bond formation was done to examine the binding stability of identified G3BP1 binding molecules. First, the structural stability was analysed using RMSD graph. This showed that the protein-ligand complexes attain stability after 10 ns and the complexes were stable throughout the simulation run up to 50 ns as shown in **Fig.3a**. The average RMSD values for G3BP1-L-7, G3BP1-WIN, G3BP1-SB2, G3BP1-NAL, G3BP1-DHD, G3BP1-GSK, and G3BP1-FLU were 0.37, 0.46, 0.39, 0.39, 0.44, 0.43 and 0.42 respectively, whereas it was 0.43 nm for ligand free G3BP1. RMSD values of all protein-ligand complexes were in the range of 0.37 and 0.46 nm which were almost comparable to that of the ligand free protein. However, all seven complexes showed comparably lower RMSD values than the ligand-free G3BP1 indicating more stable ligand-bound structures. The average fluctuation of all residues was considered and plotted as RMSF to address the residual flexibility in G3BP1 in the ligand-free state and upon ligand binding. The residual mobility of protein-ligand complex was calculated and analysed with respect to the residue number. Analysis of RMSF plot indicated that the loop regions showed higher flexibility, as depicted in **Fig.3b**, however, the secondary structural elements presented comparatively smaller deviations. The average RMSF values obtained for all the ligands fall in the range between 0.12 to 0.18 nm whereas it was 0.13 nm for the ligand-free G3BP1. Furthermore, all seven ligands did not fluctuate at the binding site, as indicated by cumulative RMSF data, defining a stable protein-ligand complex. Radius of gyration graphs of G3BP1-ligands are shown in **Fig.3c**. The conformational stability of free G3BP1 before and after binding of each ligand was investigated by calculating R_g_ of both the systems. The average R_g_ values for G3BP1 and its complexes with L-7, WIN, SB2, NAL, DHD, GSK, and FLU were 1.467nm, 1.46nm, 1.45nm, 1.47nm, 1.45nm, 1.47nm, and 1.46nm, respectively which were consistent for all the structures. The R_g_ graph suggested no major changes in the packing of G3BP1 upon binding to the ligands. This indicated that the protein-ligand structures are stable and do not affect the protein folding pattern and stability. The number and distribution of H-bonds in G3BP1- ligand complexes were analysed using g_h bond utilities of GROMACS to determine the stability of the protein-ligand complexes as shown in **Fig.3d**. A maximum number of H-bonds are formed between G3BP1-NAL (six H-bonds), followed by G3BP1-DHD (five H-bonds), then G3BP1-L-7 and G3BP1-GSK (four H-bonds), followed by G3BP1-SB2 and G3BP1-FLU (three H-bonds) and lastly, G3BP1-WIN (two H-bonds). The formation of H-bonds was consistent between the ligand-protein complexes during the complete simulation run, as shown in **Fig.3d**.

**Fig.3.**
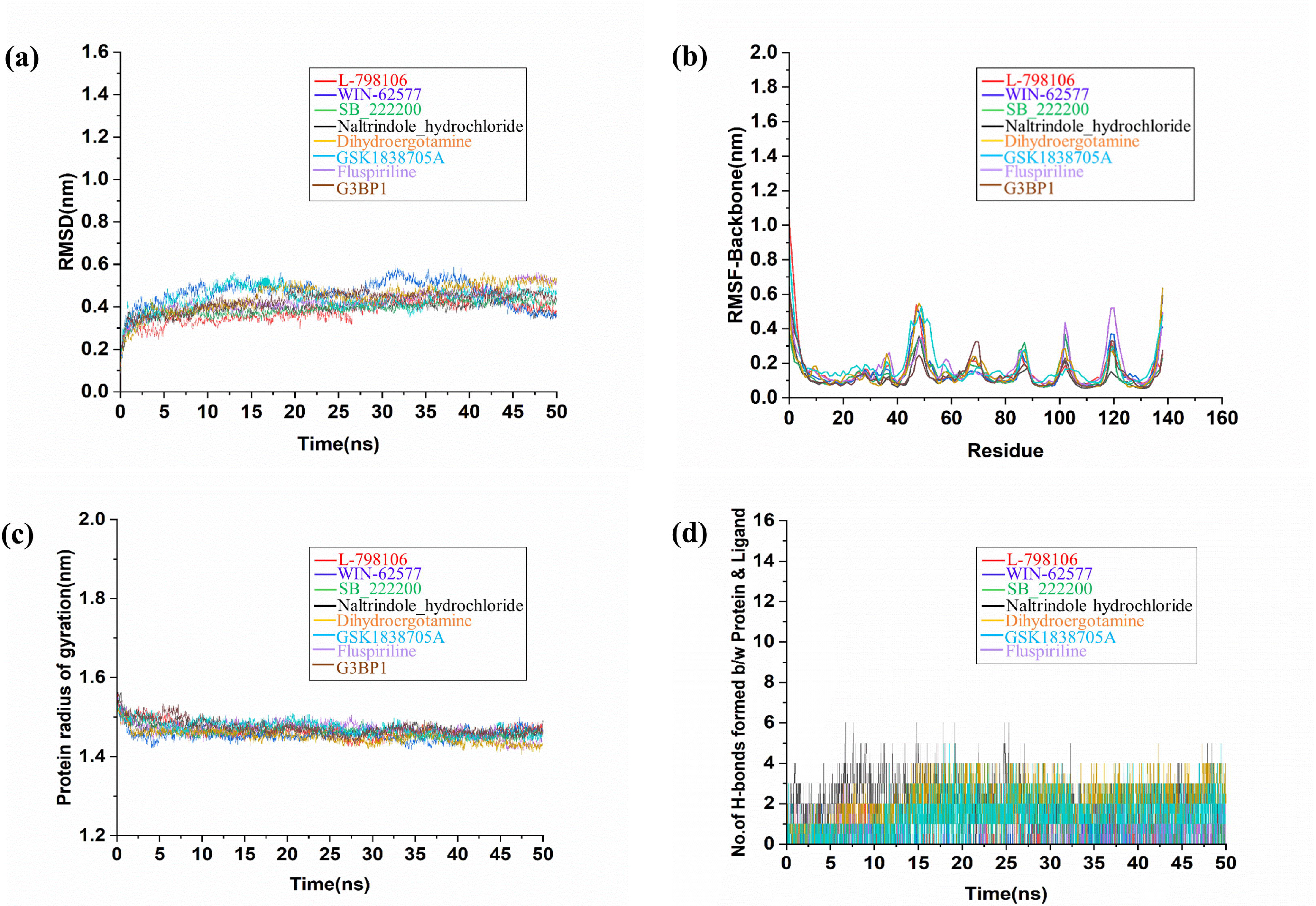
Structural dynamics and compactness of G3BP1 upon ligand binding as a function of time. **(a)** Root Mean Square Deviation (RMSD) plot of G3BP1 in complex with ligands. **(b)** Root Mean Square Fluctuation (RMSF) plot for residual fluctuations of G3BP1 before and after ligand binding. **(c)** Plot for time evolution of radius of gyration (Rg). **(d)** Hydrogen bond formation plot between G3BP1-ligand complexes. G3BP1-L-7, G3BP1-WIN, G3BP1- SB2, G3BP1-NAL, G3BP1-DHD, G3BP1-GSK, and G3BP1-FLU are represented in red, blue, green, black, orange, sky blue, light blue and brown respectively.

### Binding affinities of G3BP1 inhibitors

To evaluate the binding affinities of L-7, WIN, SB2, NAL, DHD, GSK, and FLU, ITC was performed using purified G3BP1 protein with microcalorimeter (MicroCal iTC200, Malvern, Northampton, MA) at 25 °C. The heat of dilution was negligible when separate titrations of the ligands were conducted with buffer solution in control experiments. The compounds L-7, WIN, SB2, NAL, DHD, GSK, and FLU in the syringe were titrated into the cell containing 20 µM G3BP1 protein at a concentration of 500 µM, 200 µM, 500 µM, 500 µM, 500 µM, 500 µM, 500 µM respectively by keeping all other parameters constant. Isotherms were fitted using one set of sites model for non-linear curve to conclude the thermodynamic parameters such as n, *K_D_*, ΔH, and ΔS (**Table 2**). The data was analysed using Origin7.0 associated with Microcal-ITC200 analysis software and equilibrium dissociation constant, *K_D_* values were obtained. The calculated *K*_D_ for the binding of L-7, WIN, SB2, NAL, DHD, GSK, and FLU was 18.7 μM, 58.4 μM, 66 μM, 81.3 μM, 54 μM, 98 μM, and 71.4 μM, respectively (**Fig.4**.).

**Fig.4.**
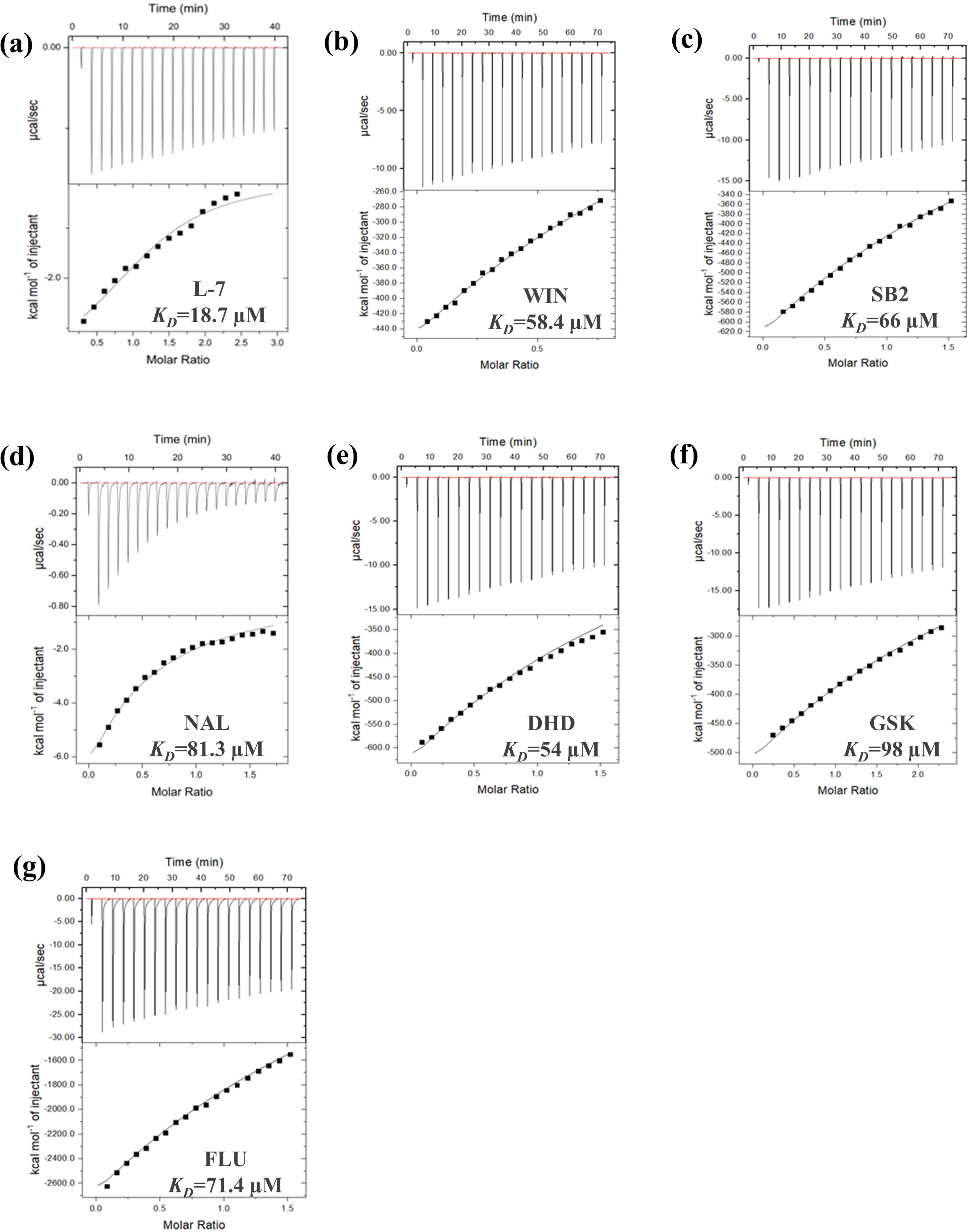
ITC profiles of molecules targeting the FGDF motif binding pocket in G3BP1. Binding isotherm for **(a)** G3BP1-L-7 Titrations. **(b)** G3BP1-WIN titrations. **(c)** G3BP1-SB2 titrations. **(d)** G3BP1-NAL Titrations. **(e)** G3BP1-DHD Titrations. **(f)** G3BP1-GSK Titrations. **(g)** G3BP1-FLU Titrations.

**Table 2.** Analysis of L-7, WIN, SB2, NAL, DHD, GSK, and FLU binding to G3BP1 as obtained from ITC.

| Protein | Ligand | n | $K_D(\mu\text{M})$ | $K_A(\text{M}^{-1})$ | $\Delta H(\text{cal/mol})$ | $\Delta S(\text{cal/mol/degree})$ |
| --- | --- | --- | --- | --- | --- | --- |
| G3BP1 | L-7 | 1 | 18.7 | $(53.3 \pm 8.5)10^3$ | $-3996 \pm 174.2$ | $-8.23$ |
| G3BP1 | WIN | 1 | 58.4 | $(1.71)10^4 \pm 872$ | $(-172.8 \pm 6.194)10^4$ | $-5.78\text{E}^3$ |
| G3BP1 | SB2 | 1 | 66 | $(1.51)10^4 \pm 578$ | $(-46.51 \pm 1.381)10^5$ | $-1.56\text{E}^4$ |
| G3BP1 | NAL | 1 | 81.3 | $(12.3 \pm 1.52)10^3$ | $-7.689 \pm 1.981$ | $-2.58\text{E}^6$ |
| G3BP1 | DHD | 1 | 54 | $(18.5 \pm 1.28)10^3$ | $(-39.27 \pm 2.050)10^5$ | $-1.31\text{E}^4$ |
| G3BP1 | GSK | 1 | 98 | $(1.02)10^4 \pm 329$ | $(-54.60 \pm 1.412)10^5$ | $-1.83\text{E}^4$ |
| G3BP1 | FLU | 1 | 71.4 | $(1.40)10^4 \pm 702$ | $(-213.5 \pm 8.503)10^5$ | $-7.15\text{E}^4$ |

Additionally, SPR biosensor is a sensitive tool to study the kinetics of ligand analyte molecular interactions as measurements of molecular binding events are done in real time. In this study, purified His-tagged G3BP1 protein was captured directly on an NTA sensor chip through Ni-mediated affinity capturing. A capture level between 800-1000 resonance units (RU) was obtained for the purified G3BP1 on injecting the protein. The binding of six compounds – L-7, WIN, SB2, NAL, DHD, GSK, and FLU was performed. For the SPR kinetic studies, the contact time of L-7, WIN, NAL and DHD binding to G3BP1 was monitored for 60s and for SB2 and FLU it was 30s, and the flow rate was set to 30 μl/min. Kinetic parameters for binding of L-7, WIN, SB2, NAL, DHD, and FLU to the G3BP1 protein were assessed. The data obtained from these experiments was evaluated by the Biacore T200 Evaluation Software, Version: 2.0. The data was fitted with one state model as per the global fitting of the kinetic model and stoichiometry 1:1(**Fig.5**). The *K_D_* values obtained from the binding of L-7, WIN, SB2, NAL, DHD, and FLU to G3BP1 from SPR were 14.34 μM, 3.4 μM, 18 μM, 46 μM, 217.2 μM and 71 μM, respectively. However, the binding constant for GSK could not be determined due to high bulk effect seen when binding kinetics SPR experiments were setup for GSK. All compounds exhibited concentration-dependent binding with G3BP1 protein.

**Fig.5.**
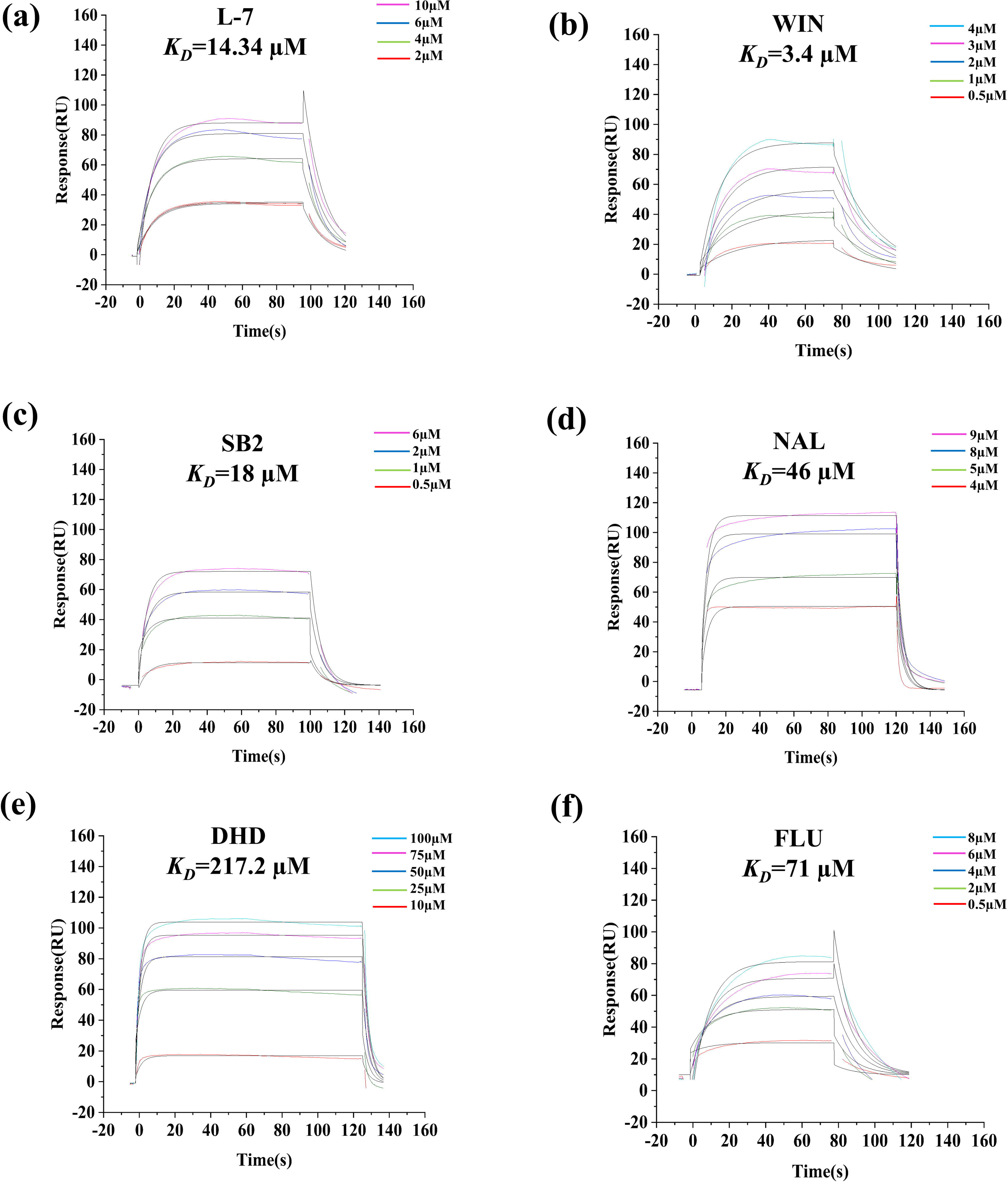
Sensograms of binding of ligands to G3BP1. (a) L-7 to G3BP1. (b) WIN to G3BP1. (c) SB2 to G3BP1. (d) NAL to G3BP1. (e) DHD to G3BP1. (f) FLU to G3BP1. Various ligand concentrations in the range of 0.5 to 100 µM were used for binding experiments to the G3BP1 protein and are shown in different colored lines.

### Evaluation of antiviral efficacy

First, the cytotoxicity assessment of the compounds was done using the MTT assay. Vero cells were treated with increasing concentrations of compounds (L-7, WIN, SB2, NAL, DHD, GSK, and FLU) for 48 h and the cell cytotoxicity was assessed. The percentage cell cytotoxicity was calculated against the untreated control. The maximal nontoxic dose (MNTD) values that showed below 10% cell cytotoxicity for L-7, WIN, SB2, NAL, DHD, GSK, and FLU were found to be 25 µM, 6.25 µM, 200 µM, 50 µM, 35 µM, 6.25 µM, and 12.5 µM respectively (**Fig.6**). The DMSO as control was also assessed with the respective concentration in the compounds solutions and was found non-toxic to Vero cells up to 1.1 % v/v concentration. Moreover, the CC_50_ value was 31.12 µM, 10.31 µM, >200 µM, 59.76 µM, 21.42 µM, 13.64 µM, and 12.47 µM for L-7, WIN, SB2, NAL, DHD, GSK, and FLU compounds, respectively.

**Fig.6.**
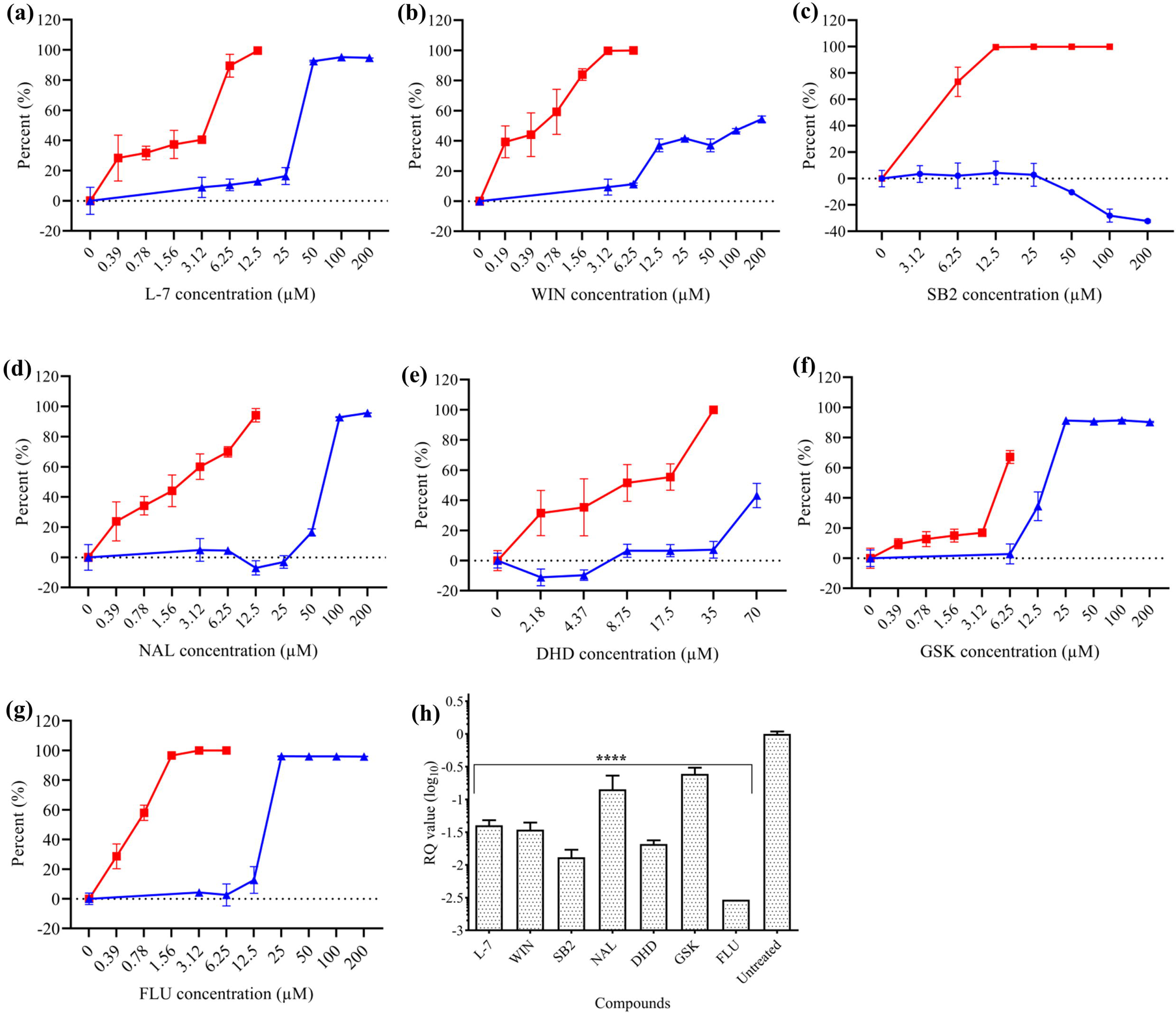
Evaluation of cell cytotoxicity and antiviral activity in cell-based assay. The percent cell cytotoxicity of Vero cells at diverse concentrations of compounds (L-7, WIN, SB2, NAL, DHD, GSK, and FLU) against the untreated control was calculated using MTT assay. Plaque reduction assay were performed with varied concentrations of compounds (L-7, WIN, SB2, NAL, DHD, GSK, and FLU) and infected cells without compounds were used as untreated control. The virus titer was calculated in pfu/mL. Then, the percent CHIKV inhibition was calculated against the untreated control and the graph was plotted using Graph Pad Prism Version 8. The percent cell cytotoxicity (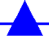) and percent CHIKV inhibition (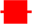) graph verse the concentration of **(a)**; compound L-7 **(b)**; compound WIN **(c)**; compound SB2 **(d)**; compound NAL **(e)**; compound DHD **(f)**; compound GSK **(g);** compound FLU. A significant reduction in infectious viral titre observed with all compounds concentration in dose dependent manner and calculated the half-maximal effective concentration (EC_50_) of all compounds (L-7, WIN, SB2, NAL, DHD, GSK, and FLU) using Graph Pad Prism Version 8. Data points represent the mean ± standard deviation of triplicate experiments. **(h) Quantification of CHIKV RNA in Vero cells by qRT-PCR:** Quantitative real-time PCR (RT- PCR) data were depicting relative quantification value (RQ=2^-ΔΔDt^) with respect to untreated virus control with 12.5 µM L-7, 6.25 µM WIN, 100 µM SB2, 12.5 µM NAL, 35 µM DHD and 6.25 µM GSK, and 6.25 µM concentration of FLU. The graphs are showing the reduction in viral RNA while plotted between RQ value (log_10_) verse compounds concentration. The experiment was done in triplicates. The statistical analysis was performed using one way – ANOVA and Dunnett’s method. Data was statistically significant as the p value is lower than 0.05 (***; p ≤ 0.001, ****; p ≤ 0.0001).

The anti-CHIKV effects of all the compounds were evaluated by performing plaque reduction assay and quantitative real time RT-PCR. Concentrations of each compound used was lower than its MNTD values to eliminate the possibility of compound mediated cytotoxicity. The viral titres were quantified using plaque reduction assay after titrating the harvested supernatants from inhibition assay at the non-cytotoxic doses. The reduction in production of infectious virus in the treatment as compared to the virus control (untreated) implied anti-CHIKV effect of compounds. Viral titre was significantly reduced in presence of the compounds (L-7, WIN, SB2, NAL, DHD, GSK, and FLU) in contrast to untreated control (**Fig.6**). Further, EC_50_ of the compounds was determined after quantification of virus particles from the plaque reduction assay. EC_50_ values were 1.996 µM ± 0.710, 0.403 µM ± 0.118, 5.387 µM ± 1.398, 1.528 µM ± 0.378, 7.394 µM ± 3.061, 3.664 µM ± 2.295, and 0.618 µM ± 0.226, for L-7, WIN, SB2, NAL, DHD, GSK, and FLU respectively, which were evaluated using non-linear regression with 95% confidence intervals using Graph Pad Prism Version 8.

To examine if the identified compounds also affects the viral genome replication, the viral mRNA gene expression levels were quantified by preforming qRT-PCR analysis. The gene expression of E1 (CHIKV specific) and actin (internal control) was quantified in real time. Relative quantification (RQ= 2^-ΔΔDt^) value showed the significant reduction in viral gene expression in treated cells for compounds (L-7, WIN, SB2, NAL, DHD, GSK, and FLU) in comparison to untreated virus control (**Fig. 6h**). Treatment with 12.5 μM L-7, 6.25 μM WIN, 100 μM SB2, 12.5 μM NAL, 35 μM DHD, 6.25 μM GSK, and 6.25 μM FLU resulted in reduction of the viral RNA level approximately by 24-fold, 29- fold, 77-fold, 7-fold, 48-fold, 4-fold, and 340-fold against the untreated virus control.

The indirect IFA was performed to reveal the effects of compound treatment on the viral load of CHIKV. IFA results showed that 12.5 µM of L-7, 6.25 µM of WIN, 100 µM of SB2, 12.5 µM of NAL, 35 µM of DHD, 6.25 µM of GSK, and 6.25 µM of FLU effectiveness in inhibiting the infectious CHIKV virus in Vero cells. The distinct reduction in the fluorescence signal of viral protein (envelope protein) was seen on treatment with compounds (L-7, WIN, SB2, NAL, DHD, GSK, and FLU) as compared to the virus control (**Fig. 7**).

**Fig.7.**
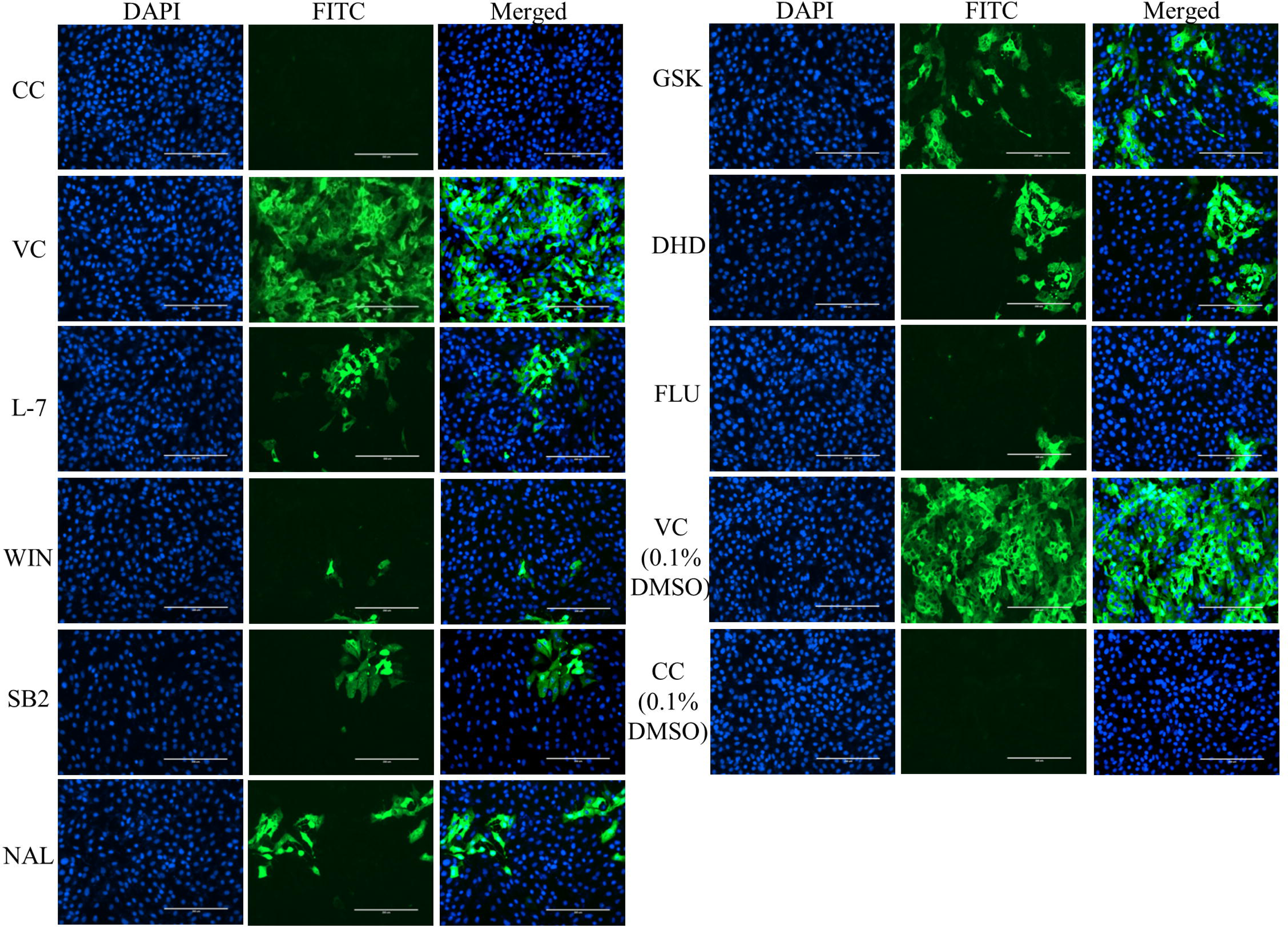
Evaluation of inhibitory effect of compounds (L-7, WIN, SB2, NAL, DHD, GSK, and FLU) treatment on CHIKV infection by Immuno-fluorescence assay. Indirect IFA of CHIKV with FITC- conjugated antibody for viral protein (envelope protein). CHIKV infected Vero cells were treated with 12.5 µM concentration of compound L-7, 6.25 µM concentration of compound WIN, 100 µM concentration of compound SB2, 12.5 µM concentration of compound NAL, 35 µM concentration of compound DHD, 6.25 µM concentration of compound GSK, and 6.25 µM concentration of compound FLU along with cell control-CC and untreated infected control-VC, both treated with 0.1 % DMSO. Cell control-CC (no virus infection and treatment), untreated infected control-VC (no treatment, only virus). Images are observed at 20X magnification and are shown for DAPI (blue), FITC (green) and Merge (blue and green).

### Formation of SGs in CHIKV infected cells and on treatment with antiviral

The formation of SGs in Vero cells was visualized using IFA. The CHIKV infected cells treated with the compounds (L-7, WIN, SB2, NAL, DHD, GSK, and FLU) showed the less virus induced SGs as compared to virus infected control. These data suggest that the compounds inhibit the CHIKV nsP3-G3BP1 interaction which reduced the CHIKV replication. If the viral replication was reduced, the viral induced SGs formation was also reduced. The distinct reduction in fluorescence of G3BP1 (SG marker protein) was seen in treatments with all compounds concentration (L-7, WIN, SB2, NAL, DHD, GSK, and FLU) as comparison to the infected control (**Fig.8**). Additionally, the infected Vero cells were evaluated under oxidative stress conditions in the presence of compounds. Identical level of SGs formation was observed in all the compounds treated infected cells as well as in the virus infected control and cell control cells (**Fig.8**). Further, the effect of compounds on the SGs formation was evaluated in the presence of oxidative stress and it indicated the compounds did not affect the assembly of SG (**Fig.8**).

**Fig.8.**
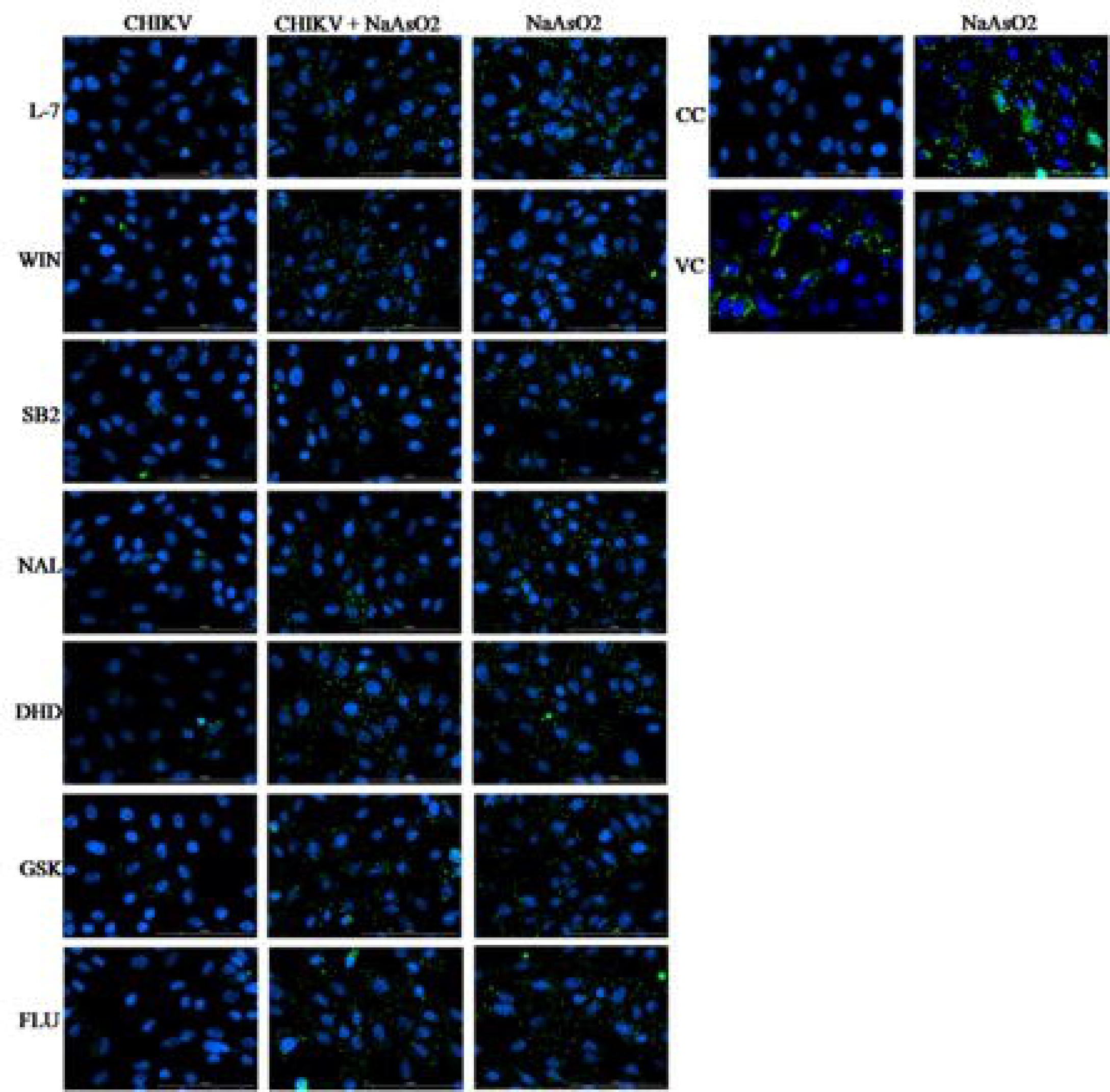
Visualization of SGs in CHIKV infected cells treated with compounds (L-7, WIN, SB2, NAL, DHD, GSK, and FLU) using immunofluorescence assay (IFA) CHIKV infected Vero cells were treated with compounds concentration (12.5 µM L-7, 6.25 µM WIN, 25 µM SB2, 50 µM NAL, 17.5 µM DHD, 3.125 µM GSK, and 3.125 µM FLU,) and then fixed at 24 hpi. Cells were stained with G3BP1-specific antibody conjugated Alexa Fluor 488 (green color), and stained with DAPI (blue color) simultaneously. Using inverted microscope, the images were captured with 40X magnification (Scale bar −100 µm). The cell controls (CC) and CHIKV infected untreated cells (VC) were kept as negative and positive controls, respectively. In the CHIKV column, the CHIKV infected cells were treated in the presence of compounds with CC and VC controls whereas in the CHIKV + NaAsO_2_ column, all cells were treated with 0.5 mM NaAsO_2_ for about 30 min. In the NaAsO_2_ column, the compounds were added in the non-infected cells and further treated with 0.5 mM NaAsO_2_ for 30 min. All images are representative examples of 3 biological repeats.

## Discussion

Viral entry triggered, G3BP1-induced SGs are antiviral structures that play an inhibitory role in the replication of several viruses. Viruses from diverse families such as *Flaviviridae*, *Paramyxoviridae*, *Caliciviridae*, *Rhabdoviridae, Togaviridae*, and others modulate SG formation in infected cells by interacting with SG effector proteins such as G3BP1. G3BPs, extensively studied marker protein of SG, have been associated in the replication of many RNA viruses [47–49]. G3BPs are also shown to interact with SINV nsP2, nsP3 and nsP4 [50–52] and SFV nsP3 [44]. CHIKV nsP3 binds G3BP1 dimers in a hierarchical manner which is critical for virus replication. It has been reported that the interaction between the NTF-2 like domain of G3BP1 and FGDF motifs in the C-terminus of viral nsP3 is necessary to recruit G3BP1 into viral replication complexes. This recruitment by nsP3 results in the disassembly of SGs [53]. Since there is less genetic variability of the host as compared to viruses, host-based antiviral therapeutics is an efficient alternative approach against chronic viral infections. Targeting the interaction between host protein G3BP1 and CHIKV nsP3 could prove to be an effective host-directed strategy for development of effective antivirals to curb the resurgence of chikungunya disease. Reportedly, in SFV, a 1.9 Å co-crystallized structure of NTF2 domain of G3BP1 with a 25 amino acid peptide was determined (PDB ID:5FW5). The revealed structure emphasizes that FGDF motifs bind into the hydrophobic groove present in the NTF2-like domain of G3BP1 which was shown to be critical for CHIKV replication. By identifying the potential small inhibitor molecules which can inhibit this G3BP1-CHIKV nsP3 protein interaction could be a promising host-directed antiviral therapy. In our study, for the first time, we have identified and proposed the small molecules endowed with antiviral activity against CHIKV targeting this host-viral protein interaction. A library of bioactive small molecules was screened *in silico* using a structure-based approach against the available crystal structure of NTF-2 like domain of G3BP1 in complex with 25 amino acid residue peptide (PDB ID:5FW5) [46]. Seven compounds based on highest binding affinity were selected (**Fig.1a-1g**). These hit compounds were further chosen for docking studies to evaluate their binding efficiency against G3BP1. Molecular docking revealed the binding profiles and molecular interactions of proposed compounds with the active site and substrate binding residues of G3BP1 **(Fig.2a-2g)**. Furthermore, dynamic simulation analysis of the G3BP1-ligand complex from the trajectory of 50ns provided us with an insight of strong binding affinity and, thus, stability (**Fig.3a-3d**). Additionally, the thermodynamic parameters of the ligand binding were calculated using ITC. The binding constant (*K*_D_) calculated for the binding of L-7, WIN, SB2, NAL, DHD, GSK, and FLU were 18.7 μM, 58.4 μM, 66 μM, 81.3 μM, 54 μM, 98 μM, and 71.4 μM, respectively **(Fig.4a-4g).** The results were further validated using an alternative biophysical technique SPR. The *K_D_* value of L-7, WIN, SB2, NAL, DHD, and FLU to purified G3BP1 as obtained in SPR kinetics was 14.34 μM, 3.4 μM, 18 μM, 46 μM, 217.2 μM and 71 μM, respectively **(Fig.5a-5f)**. Hence, *K_D_* values in micromolar range in both ITC and SPR-based studies indicate good binding affinity of the identified compounds with G3BP1 protein. The difference in the values of rate constants obtained in ITC and SPR could be due to difference types of buffers used in the experiments, parameters that are set for the experiments and different principle of these two biophysical techniques. Concordantly, the binding potential of all the compounds with G3BP1 in combination with dynamic simulation studies, ITC and SPR experiments suggest that all molecules can serve as a noble chemical scaffold for developing new antivirals against CHIKV, targeting the host-viral protein interactions.

To further strengthen our findings until now, we attempted to characterise the antiviral activity of all the compounds in *in vitro* cell-culture based antiviral assays. Prior to estimation of antiviral properties, we determined the cytotoxic effects of the compounds. The cytotoxic effects of L-7, WIN, SB2, NAL, DHD, GSK, and FLU on Vero cells were analysed by MTT assay. Cell viability assay revealed that MNTD value of L-7, WIN, SB2, NAL, DHD, GSK, and FLU was approximately 25 µM, 6.25 µM, 200 µM, 50 µM, 35 µM, 6.25 µM, and 12.5 µM respectively that showed below 10 % cell cytotoxicity at 48 h **(Fig.6a-6g)**. The CC_50_ value for L-7, WIN, SB2, NAL, DHD, GSK, and FLU was 31.12 µM, 10.31 µM, >200 µM, 59.76 µM, 21.42 µM, 13.64 µM, and 12.47 µM respectively. The antiviral activity of the compounds was evaluated at concentrations lower than their MNTD values so that the possibility of compound mediated cytotoxicity can be eliminated. Antiviral assay scanning different concentrations of all the compounds was performed to test their antiviral potential. Plaque reduction assay, qRT-PCR, and IFA assays were performed to evaluate the anti-CHIKV effects of the compounds. The viral titres were quantified using plaque reduction assay at the non-cytotoxic doses which indicated a significant reduction in viral titre with the compounds **(Fig.6a-6g)**. The reduction in production of the infectious virus in the test experiments implied anti-CHIKV effect of the compounds. EC_50_ values, thus obtained, after plaque reduction assay were 1.996 µM ± 0.710, 0.403 µM ± 0.118, 5.387 µM ± 1.398, 1.528 µM ± 0.378, 7.394 µM ± 3.061, 3.664 µM ± 2.295, 0.618 µM ± 0.226 for L-7, WIN, SB2, NAL, DHD, GSK, and FLU respectively. Furthermore, the data obtained from qRT-PCR confirmed the decrease in CHIKV RNA expression in the cells treated with the compounds after 24 h, hence, suggested a decrease in CHIKV replication. We observed approximately 24-fold, 29-fold, 77-fold, 7-fold, 48-fold, 4-fold, and 340-fold reduction in CHIKV RNA expression levels as compared to untreated virus control when the Vero cells were treated with 12.5 μM L-7, 6.25 μM WIN, 100 μM SB2, 12.5 μM NAL, 35 μM DHD, 6.25 μM GSK and 6.25 μM FLU, respectively (**Fig. 6h**). Altogether, we can postulate that the compounds are inhibiting the G3BP1- nsP3 interaction, thereby, substantially lowering the CHIKV replication. Additionally, IFA experiment was conducted to assess the anti-CHIKV activity of the compounds. The decrease in fluorescence of viral protein was observed in infected cells treated with compounds as compared to infected control. IFA results further revealed the anti-CHIKV potential of the compounds (**Fig.7**).

To investigate the relationship between CHIKV replication and SG formation, we proceeded with IFA experiment to visualize the formation of SGs in Vero cells. Earlier, it was reported that replication of CHIKV replicon induce G3BP1-containing SGs, and nsP3 expression alone was sufficient to sequester G3BP1 into granules [45]. We also established that the CHIKV infected cells treated with these compounds (L-7, WIN, SB2, NAL, DHD, GSK, and FLU) showed less virus-induced stress granules as compared to virus infected control. It reinforced that these compounds inhibit CHIKV nsP3-G3BP1 interaction which contributes to reduction in CHIKV replication. We also report the distinct reduction in fluorescence signal of G3BP1 (SG marker protein) in virus-infected cells treated with compounds as compared to infected control. This supports the fact that the reduced viral replication is responsible for decrease in virus–induced stress granules (**Fig.8**). In addition, we analysed the effect of the compounds in the presence of oxidative stress given by sodium arsenite treatment. However, no effect on SG assembly was reported (**Fig.8**).

In summary, our approach of conjoining *in silico* workflow, biophysical kinetic studies, and *in vitro* studies for inhibitors of CHIKV nsP3-G3BP1 interaction identified seven novel compounds as effective inhibitors that reduced viral replication in Vero cells.

## Conclusion

In response to many viral infections, the formation of cytoplasmic SGs is induced to halt the replication of virus. This is very similar to the formation of SGs during cellular stress such as hypoxia, heat, and oxidative conditions. The amalgamation of data obtained from structure-based *in-silico* computational analysis, biophysical binding validation and *in vitro* cell-culture based antiviral assays ominously supports that the compounds bind to G3BP1 and effectively curbs the viral infection. Additionally, the FGDF binding pocket of G3BP1 protein targeted in this study for identification of anti-CHIKV molecule, has been recognized to be involved in molecular interactions with other viral and stress granule components during cellular stress conditions in various infectious and non-infectious diseases. Thus, the finding of G3BP1 inhibitors in this study are expected to have a broad impact towards the development of treatments, not only for viral infections but also on various other non-infectious diseases like cancer and neurodegenerative disorders.

## Materials and methods

### Virtual screening and molecular docking of G3BP1 protein

The crystal structure of the NTF2-like domain of G3BP1 in complex with a 25-mer peptide from SFV nsP3 was retrieved from RCSB-PDB (ID: 5FW5) for structure-based *in silico* virtual screening and molecular docking studies. PyRx 0.8 [54] and AutoDock 4.2 [55] were used to perform virtual screening and molecular docking studies. The LOPAC^1280^ (library of pharmacologically active compounds) library was used to screen the top hit compounds. Virtual screening and docking were conducted using Mac Pro, 3GHz 8-core Intel Xeon E5. All the ligands were in SDF format that were first converted into Autodock ligands (.pdbqt) format and subsequently converted in minimized energy state. The grid parameters were set, grid box dimensions were set to 76 Å×68 Å×120Å and other screening parameters were set as default. Observing the binding energy (B.E.) of G3BP1-FGDF complex as reference, the molecules having higher B.E. than G3BP1-FGDF complex were selected for detailed docking and simulation studies. During docking procedure, water molecules were removed. Kollman charge (1.248) and non-polar hydrogens were added into the protein structure using AutoDock MGL Tools [56]. Then, the protein was saved in .pdbqt file format. The .pdbqt file format was retrieved from PyRx directory for the top hit compounds (ligands). The grid parameters were kept same as screening. Gasteiger charges and hydrogen atoms were added to the ligands and saved into .pdbqt file format. The atomic potential grid map was calculated by AutoGrid 4 with a spacing of 0.316 Å and grid box dimensions were same as screening *ie* (76 Å×68 Å×120Å). The centre point co-ordinates set was X= −4.173, Y= −27.601 and Z= - 14.535. For docking procedure, the combination of Lamarckian Genetic Algorithm (GA) with grid-based energy evaluation method was used. The number of total GA runs were increased to 50 with maximum number of evaluations (250,000,000). Other docking parameters were set as default. The best fit conformation of compounds was carefully selected based on their interactions with G3BP1, lowest RMSD values, and binding energies. The analysis of the protein-ligand complexes was performed using PyMOL [57] and LigPlot^+^ [58].

### Molecular dynamics simulations

From the binding energies, lowest root mean square deviation (RMSD) and details of molecular interactions of ligands, the best-docked poses were selected and subjected to molecular dynamics (MD) simulation to assess the dynamics, complex stability and structural changes at the atomistic level in the protein. MD simulation was done using Linux based GROMACS version 5.9 [59] installed on a graphical processing unit (GPU). The topology and GROMACS file for the molecules and protein were generated using PRODRG webserver [60] and GROMOS 43a1 force field respectively. Solvation of the protein was done using the simple point charge water model in a box of cubic dimensions and radius 1.5 nm. Na^+^ and Cl^−^ ions were supplemented to neutralise the system. Using the steepest descents algorithm, the system was minimized in 50000 steps to lower the steric clashes in the system. The minimized system was then subjected to equilibration using NVT (a constant number of particles, volume and temperature at 300 K) and NPT (a constant number of particles, pressure and temperature at 1 bar pressure). The entire simulation was executed for 50 ns on the system along with the data generation at every 2 fs. The in-built tools of GROMACS such as RMSD, root mean square fluctuation (RMSF), Radius of gyration (Rg) and Hydrogen bond (H-bond) were used to analyse the trajectory obtained from the protein-ligand complexes simulation run.

### Biophysical binding analysis

#### Binding affinity measurement using isothermal titration calorimetry

To assess the binding affinities between G3BP1 protein-ligand complexes, isothermal titration calorimetry (ITC) experiments were conducted using microcalorimeter (MicroCal iTC200, Malvern, Northampton, MA) [61] to conclude the thermodynamic parameters of binding of compounds to purified G3BP1 protein. G3BP1-NTF2 (residues 1-139) protein with a His-tag was expressed and purified using immobilized metal affinity chromatography as reported earlier [46]. His-tag was cleaved using TEV protease and reverse nickel affinity chromatography was performed to remove His-tagged protein as well as His-tagged TEV protease. Buffer exchange of the purified protein was done from Tris buffer to 1X PBS buffer and was concentrated up to 5.5 mg/ml. During the experiments, seven ligands (**Table 3**) in the range of 200-500 µM were filled in the syringe and titrated into 20 µM of purified G3BP1 protein contained in the cell of ITC instrument. After equilibration, a total of 20 injections of syringe samples were titrated in the cell with one injection of 0.4 μL followed by 19 injections of 2 μL at 220 s of interval. The parameters for the experiments were set to a stirring rate of 800 rpm within the cell, a constant temperature of 25 °C and a reference power of 9 µcal/s. The ITC data was integrated, dilution effects were corrected, and concentration was normalized before the curve fitted to a one site binding model using Malvern’s Origin 7.0 Microcal-iTC200 analysis software. Moreover, reaction stoichiometry (n), the dissociation constant (K*_D_*), c value (c) as well as the thermodynamic parameters such as entropy (ΔS), enthalpy (ΔH), and Gibbs free energy change (ΔG) were measured to estimate the binding strength of all the seven molecules with the protein target.

**Table 3.** Details of all seven compounds, including stock solution and solubility are summarized.

| S.No. | Compounds Details | Catalog no. | Stock solution and Solubility | Symbol |
| --- | --- | --- | --- | --- |
| 1 | Naltrindole_hydrochloride | #N115-5MG (Sigma-Aldrich) | 8 mg/mL in Water (17.74 mM) | NAL |
| 2 | L-798106 | #L4545-5MG (Sigma-Aldrich) | 20 mg/mL in DMSO (37.28 mM) | L-7 |
| 3 | WIN_62577 | #W104-5MG (Sigma-Aldrich) | 4 mg/mL in DMSO (9.14 mM) | WIN |
| 4 | SB_222200 | #S5192-10MG (Sigma-Aldrich) | 28 mg/mL in DMSO (73.6 mM) | SB2 |
| 5 | Fluspiriline | #19530-5MG (Cayman) | 20 mg/mL in DMSO (21 mM) | FLU |
|  |  | Chemical) |  |  |
| 6 | Dihydroergotamine | #23847-25MG<br>(Cayman<br>Chemical) | 20 mg/mL in<br>DMSO (29.4 mM) | DHD |
| 7 | GSK1838705A | #SML0995-5MG<br>(Sigma-Aldrich) | 20 mg/mL in<br>DMSO (37.6 mM) | GSK |

#### Surface plasmon resonance

SPR studies for studying binding kinetics were carried out using BIACORE T200 (GE Healthcare, USA) with Ni-NTA research grade sensor chip (GE Healthcare, USA). The temperature was maintained at 25°C for analysis and sample compartments for all the binding and kinetic experiments. The SPR experiments were carried out in 1X PBS (GE-healthcare) with 0.2 %Tween 20 (catalog no: P9416-Molecular Biology grade). Dilutions of the analyte stock, samples, and buffer blanks were prepared in 1X PBS. Distilled deionized water was used for preparing all buffers. To avoid microbubbles formation, water was filtered using Millipore filters 0.22 mm and degassed using Milli-pore degassing unit. The purified His-tagged NTF2-like domain of G3BP1 was directly captured on an NTA sensor chip via Ni-mediated affinity capturing. An injection of 0.5 mM NiCl_2_ for 1 min was injected followed by purified G3BP1 protein at a concentration of 20 µM (0.4 mg/ml) with a flow rate of 30 μl/min and contact time of 120 s for the direct NTA chip capture. Flow cell 3 (FC-3) was selected as a reference cell to minimize the unwanted drift and systematic noise. At the end of each binding cycle, the G3BP1 surface was regenerated with injection of 350 mM EDTA for 45 s at the flow rate of 30 µl/min. The constant flow rate was maintained throughout the binding (30 μl/min) and kinetics experiment (30 μl/min). All the ligand samples at a concentration range from 0.5–100 µM were injected over 30-60 s period at a flow rate of 30 µl/min to study binding kinetics. All the ligands were tested in duplicates for binding to G3BP1 in PBS-P^+^ running buffer, which states that the activity of G3BP1 was retained throughout the experiment. The SPR data was collected using Biacore control software and analyzed with T200 evaluation software.

#### Cells, virus and compounds

Vero cell line was procured from National Centre for Cell Science (NCCS), Pune, India and maintained in Dulbecco’s Modified Eagle Medium (DMEM) supplemented with 10 % heat inactivated Fetal Bovine Serum (FBS) and 1x Pen-Strep (antibiotics solution). Temperature and humidity conditions were adjusted to 37 °C with 5 % CO_2_ and 95 % respectively. For this study, a clinical isolated strain of CHIKV was propagated in Vero cells [62]. DMEM (high glucose), Minimum Essential medium Eagle (MEM), Sodium Bicarbonate, Dulbecco’s Phosphate Buffered Saline (PBS), Trypsin- EDTA solution, MTT [3-(4,5-dimethylthiazol-2- yl)-2,5-diphenyltetrazolium bromide], Dimethyl Sulfoxide (DMSO), and Formaldehyde (37 %) were obtained from HiMedia, India. FBS, Pen-Strep (10,000 units/mL of penicillin and 10,000 µg/mL of streptomycin) was acquired from Gibco, USA. Details of all the seven compounds including stock solution and solubility are summarized in **Table 3**. The working concentrations for each compound (L-7, WIN, SB2, NAL, DHD, FLU and GSK) for doing antiviral assays were diluted and prepared using cell culture medium.

#### Cytotoxicity of compounds

A colorimetric MTT assay was performed to measure the cytotoxic effects of identified compounds before testing the antiviral activity. The Vero cells were seeded onto 96-well culture plate at optimum cell density (2*10^4^ cells/mL) per well. Upon reaching the 90 % monolayer confluency, the medium was removed and different concentrations of the compounds (L-7, WIN, SB2, NAL, DHD, FLU and GSK) (200 µM, 100 µM, 50 µM, 25 µM, 12.5 µM, 6.25 µM and 3.12 µM of L-7, WIN, SB2, NAL, and FLU; 70 µM, 35 µM, 17.5 µM, 8.75 µM, 4.37 µM and 2.18 µM of DHD; and 200 µM, 100 µM, 50 µM, 25 µM, 12.5 µM and 6.25 µM of GSK) diluted in DMEM with 2 % FBS were added to respective wells along with the untreated control without compounds. For cytotoxic effects of dimethyl sulfoxide (DMSO), the cells were treated with different concentration of DMSO (2.2, 1.1, 0.55, 0.27 and 0.13 % v/v) only. After 48 h of incubation, MTT (5 mg/mL) was added in each well and incubated in dark at 37 °C for 4 h. The medium containing MTT was aspirated, and the formed formazan crystals were dissolved in DMSO. The reading was taken at 570 nm using multimode-plate reader (Synergy, BioTek Instrument INC). Each experiment was done in triplicates. The percent cell cytotoxicity was calculated against the untreated control and the half maximal cytotoxic concentration (CC_50_) value was calculated with the help of Graph Pad prism Version 8 for all seven compounds.

#### Inhibition assay

Vero cells at 2×10^5^ cells/well cell density were seeded in 24-well cell culture plate. After obtaining the optimum confluency, Vero cells were treated with the different concentrations of compounds (12.5 µM to 0.39 µM of L-7; 6.25 µM to 0.19 µM of WIN; 100 µM to 6.25 µM of SB2; 12.5 µM to 0.39 µM of NAL; 35 µM to 2.18 µM of DHD; 6.25 µM to 0.39 µM of GSK; and 6.25 µM to 0.39 µM of FLU) with untreated virus control (without compound) diluted in DMEM with 2 % FBS for cells 12 h prior to CHIKV infection. Cells treated with compounds were then infected with CHIKV at multiplicity of infection (MOI) of 1 for 90 min in presence of the compounds with the same concentrations of the respective wells. After infection, the virus inoculum was removed, and the excess virus was removed from the infected cells by washing. DMEM with 2 % FBS containing the compounds was then added as the post-infection media into the respective wells for 24 h. At 24-hour post infection (hpi), supernatant from each well was harvested and the viral titre was quantified using plaque reduction assay. The cells were also harvested for determining the reduction in viral RNA due to the treatment of cells with compounds by real time reverse transcription-polymerase chain reaction (RT-PCR).

#### Plaque reduction assay

Vero cells were seeded into 24-well culture plates at desired cell density. At optimum confluency (more than 90 %), the serial dilutions of the supernatants harvested after treatment with compounds in the inhibition assay were inoculated onto cells for 90 min. Gentle shaking was done frequently and after the adsorption of virus for 90 min, the inoculum was removed. The cells were overlaid with 1:1 volume of 2 % methyl cellulose (Sigma, USA) in 2X MEM supplemented with 5 % FBS and plate was incubated for 48 h. After discarding the overlay media, the cells were fixed with 10 % v/v formaldehyde and stained with 1 % w/v crystal violet for plaque visualisation. The plaques were counted for determination of the titer in pfu/mL. The percent CHIKV inhibition was then calculated against the untreated control. All the experiments were performed in triplicate. The graphs were plotted using Graph Pad Prism Version 8.

Further, dose-dependent response curves were plotted by calculating pfu/ml in plaque assay for different compound concentrations. A range of diverse concentrations of each compound (L-7, WIN, SB2, NAL, DHD, GSK, and FLU) were used to generate inhibition curves for calculating their half maximal effective concentration (EC_50_) from generated inhibition curves. The EC_50_ value of each compound was assessed using non-linear regression with 95 % confidence intervals with the help of Graph Pad Prism Version 8. Data points indicate the mean ± standard deviation of triplicate experiments.

#### Quantitative real-time RT-PCR (qRT-PCR)

The reduction in viral RNA levels was determined using qRT-PCR from the supernatant of inhibition assay with 12.5 µM concentration of L-7, 6.25 µM concentration of WIN, 100 µM concentration of SB2, 12.5 µM concentration of NAL, 35 µM concentration of DHD, 6.25 µM concentration of GSK, and 6.25 µM concentration of FLU and untreated virus control. The total RNA was isolated from 24 h post infected cells using TRI Reagent® (Sigma-Aldrich) according to the protocol described by the manufacturer. One Micro volume UV-Vis spectrophotometer (Thermo Scientific™). Using the PrimeScript first strand cDNA synthesis kit (Takara, cat no- #6110A), the first strand of cDNA was synthesised according to the manufacturer’s instructions, using an equal amount of random primers and RNA (30 ng) for each sample to perform reverse transcription reaction. The cDNA was subsequently used to amplify CHIKV E1 coding region along with beta-actin (internal control) using KAPA SYBR® FAST qPCR master mix (2X) kit and following primer sequences were used for E1; forward: 5’- AAGTACACTGTGCAGCTGAGT-3’ and reverse: 5’- GCATAGCACCACGATTAGAATC-3’ and for actin; forward: 5’- ATTGCCGACAGGATGCAGAA-3’[63]. The amplification was performed using Quant Studio 5 Real Time PCR system (Applied Biosystems). To calculate the relative quantification value for each compound, ΔΔCt method (RQ = 2^-ΔΔDt^) was used and RQ was compared relative to CHIKV infected control (untreated). The experiments were performed in triplicate. The statistical analysis was done by one way – ANOVA and Dunnett’s method of Prism® 8 software (GraphPad Software, San Diego, CA).

#### Indirect immunofluorescence assay (IFA)

Vero cells at a density of 2×10^5^ cells/well were seeded in 24-well cell culture plate. At optimum confluency, cells were treated with 12.5 µM of L-7, 6.25 µM of WIN, 100 µM of SB2, 12.5 µM of NAL, 35 µM of DHD, 6.25 µM of GSK, and 6.25 µM of FLU, with virus infected control (VC- without compound) and cell control (CC- without virus and compounds treatment). These were diluted in DMEM with 2 % FBS for cells 12 h prior to CHIKV infection. The compound-treated cells were then infected with CHIKV at MOI of 1 for 90 min along with the same compound concentrations to respective wells. After infection, the virus inoculum was removed, and the excess virus from the infected cells was removed by washing the cells. DMEM with 2 % FBS containing the same concentration of these compounds was added into respective wells, as the post-infection media for 36 h. Similarly, virus infected control (VC) and cell control (CC) treated with 0.1 % DMSO were also taken as the controls for this assay. At 36 h post infection (hpi), the cells were rinsed with chilled PBS. The fixing reagent [methanol: acetone (1:1)] was added for 1 h at room temperature (RT). 0.1 % Triton X-100 (Qualigens, India) added for 7-10 min for permeabilization. The cells were washed three times with PBS. The cells were then incubated with primary antibody, anti-alphavirus monoclonal antibody (mAb) (Santa Cruz Biotechnology, USA, cat. no. sc-58088) (1:100) in PBS for 1-2 h. After washing the cells thrice with PBS, the cells were incubated with secondary antibody in PBS (1:250) anti-Mouse IgG (whole molecule) –fluorescein isothiocyanate (FITC) antibody produced in goat (Sigma, F9006) in dark for 1 h. After washing thrice with PBS, nuclei were stained with 4′,6-diamidino-2-phenylindole (DAPI) (20 µg/mL). Cell imaging using EVOS FL Auto Imaging System (Thermo Fisher Scientific) was done at 20X magnification.

#### IFA for visualization of SGs

Indirect IFA was performed for visualization of SGs in the treated and untreated CHIKV infected cells. Vero cells at a cell density of 3×10^5^ cells/mL were seeded on glass coverslips in 6-well plates overnight. Then, media was replaced with media containing compounds at the non-cytotoxic concentration (12.5 µM L-7, 6.25 µM WIN, 25 µM SB2, 12.5 µM NAL, 17.5 µM DHD, 3.12 µM GSK and 3.125 µM FLU) with virus infected control (VC- without compound) and cell control (CC- without virus and compounds treatment), both diluted in DMEM with 2 % FBS for cells 8 h prior to CHIKV infection. The cells were then infected with CHIKV at MOI of 1 for 90 min along with the same compounds concentration to respective wells was done. After infection, cells were washed to remove the excess virus and DMEM with 2 % FBS containing the identical concentration of the compounds was added into respective wells. Similarly, cells treated with compounds only were also taken as the controls for this assay to visualize the effect of compounds on SGs assembly. Cells were incubated for 24 h at 37 °C before induction of stress with 0.5 mM sodium arsenite (NaAsO_2_) 30 min prior to fixation by 3.7 % formaldehyde. The fixed cells were permeabilized using methanol for about 10 min at −20 °C, and blocked using 3 % bovine serum albumin (BSA) in PBS for 1 h. Then, immersed in 1:100 dilution of G3BP1 (H-10), mouse monoclonal antibodies tagged Alexa Fluor 488 (sc-365338 AF488) at 4 °C for overnight and nuclei were stained using DAPI. Following washes with PBS, cover slips containing cells were mounted using VITROCULD mounting media (Deltalab) and viewed using Lionheart LX Automated Microscope (Agilent BioTek) at 40X magnification. Images were captured and processed for adjustments in brightness and contrast, color, exposure etc and were saved as TIFF files.

## Acknowledgement

Authors, SM, and AS acknowledge to Ministry of Human Resource Development, Government of India for financial support and AP acknowledge the Council of Scientific & Industrial Research (CSIR) for providing research fellowship. RK acknowledge to Ministry of Human Resource Development, Government of India for financial support for Post- Doctoral Fellowship. PK and ST thank Department of Biotechnology, Govt of India for supporting Bioinformatics Center at IIT Roorkee (reference number BT/PR40141/BTIS/137/16/2021). PK thanks to STARS, MOE (Project ref no: STARS2/2023-0209) for funding this study. Authors express special thanks to Macromolecular Crystallographic Unit (MCU), Institute instrumentation centre (IIC) at Indian Institute of Technology (IIT) Roorkee for providing the facility to carry out molecular simulation studies. Authors thank the MCU at Institute Instrumentation Centre, Indian Institute of Technology (IIT) Roorkee for providing facility to carry out ITC experiments. Authors express special thanks to Prof. Debrupa Lahiri and her research scholar Mr. Arabinda Majhi, 3D Bio-printing and cell-culture facility, RETHINK-Tinkering laboratory, IIT Roorkee for providing inverted fluorescence microscope facility to preform fluorescence imaging.

## Author contributions

S.T., P.K., and S.M. conceptualized; S.M., R.K., A.S., and A.P. applied the methodologies; S.T. investigated the methodologies; S.M. and R.K. performed the formal analysis and analysed the data; S.M. prepared the original draft; S.M. and R.K. and S.T. edited the manuscript; S.M., R.K. and A.S. prepared the figures; P.K.provided some resources; S.T. reviewed and supervised the manuscript.

